# Novel gut pathobionts confound results in a widely used mouse model of human inflammatory disease

**DOI:** 10.1101/2021.02.09.430393

**Authors:** Samuel C. Forster, Simon Clare, Benjamin S. Beresford-Jones, Katherine Harcourt, George Notley, Mark Stares, Nitin Kumar, Amelia T. Soderholm, Anne Adoum, Hannah Wong, Bélen Morón, Cordelia Brandt, Gordon Dougan, David J. Adams, Kevin J. Maloy, Virginia A. Pedicord, Trevor D. Lawley

**Affiliations:** Host-Microbiota Interactions Lab, Wellcome Sanger Institute, Hinxton, United Kingdom; Centre for Innate Immunity and Infectious Diseases, Hudson Institute of Medical Research, Clayton, Victoria, Australia; Department of Molecular and Translational Sciences, Monash University, Clayton, Victoria, Australia; Cambridge Institute of Therapeutic Immunology and Infectious Disease, Jeffrey Cheah Biomedical Centre, Cambridge, United Kingdom; Department of Medicine, University of Cambridge School of Clinical Medicine, Cambridge, United Kingdom; Animal Health Trust, Lanwades Park, Kentford, Newmarket, Suffolk, United Kingdom; Institute of Infection, Immunity and Inflammation, University of Glasgow, Glasgow, United Kingdom

**Author notes:** corresponding author Trevor D. Lawley, Wellcome Sanger Institute, Hinxton, Cambridgeshire, UK.

**Keywords:** microbiota, pathobiont, inflammatory bowel disease, dextran sulfate sodium

## Abstract

The mammalian gut microbiota consists of hundreds of anaerobic bacterial species that shape intestinal homeostasis and influence host immune responses. Although the causal roles of specific human gut bacterial species in health and disease are emerging, the role of indigenous gut bacteria in driving immunophenotypic variability in mouse models of human disease remains poorly understood. We performed a large-scale experiment using 579 laboratory mice designed to identify and validate the causes of disease variability in the widely used dextran sulphate sodium (DSS) mouse model of inflammatory bowel disease. Using microbiome analysis, coupled with machine learning and targeted anaerobic culturing, we identified and isolated the novel gut pathobiont species *Duncaniella muricolitica* and *Alistipes okayasuensis* and fulfilled Koch’s postulates in mice to show that each pathobiont exerts dominant effects in the DSS model leading to variable treatment responses. We show these pathobiont species are common, but not ubiquitous, in mouse facilities around the world, raising experimental design opportunities for improved mouse models of human intestinal diseases.

---

Experimental mouse models are central to basic biomedical research and serve as important pre-clinical models in drug discovery for a variety of human diseases, including autoimmune and metabolic disorders, cancers and infections. Unfortunately, there is tremendous variability in disease penetrance and reproducibility among genetically identical mice within and between mouse facilities, limiting the utility of many commonly used models^1-3^. Specific pathogen free (SPF) protocols have been successfully employed to control the confounding effects that can be caused by known bacterial pathogens, but we lack equivalent knowledge of the effects of the diverse, uncharacterized symbiotic bacteria within the indigenous gut microbiota of mice. While our knowledge of the human gut microbiota in health and disease has expanded extensively with large-scale microbiome studies, comprehensive computational bacterial discovery^4,5^, and extensive genome sequenced bacterial collections^6-9^, equivalent investigations in mouse models has remained limited. The vast differences in species type, composition and dispersal between human and mouse microbiotas highlight the importance of efforts to culture mouse specific bacteria and to validate their functions during intestinal homeostasis and disease^10,11^.

Here we describe a large-scale experiment using 579 genetically identical laboratory mice designed to identify and validate the causes of variability for standard experimental endpoints (weight loss and intestinal pathology) in the widely used Dextran Sulphate Sodium (DSS) mouse model of inflammatory bowel disease (IBD)^12^. Using a data-driven approach, we identified from gut microbiome data, candidate bacterial taxa driving weight loss and intestinal inflammation. We then cultured and functionally validated that these specific, novel bacterial strains can alone determine experimental outcomes in gnotobiotic mice. Thus, using a Koch’s postulate approach for microbiome discovery, we identified the novel mouse gut pathobiont species, *Duncaniella muricolitica* and *Alistipes okayasuensis*, found frequently in mouse colonies around the world, that have a dominant impact on DSS model outcomes.

## RESULTS

### Disease severity is highly variable in the DSS mouse model

Inflammatory bowel disease (IBD) research relies heavily on laboratory mice to understand the role of host genetics in disease resistance and susceptibility; however, the gut microbiota is rarely profiled at a genomic level or cultured in the lab, nor robustly controlled to understand the contribution of gut bacteria to experimental outcomes. Potential modulators of IBD are routinely assessed in mice through administration of DSS in drinking water^12,13^, which acts by damaging the gut epithelial barrier to trigger inflammatory disease. To quantify the variability of the DSS model and to investigate primary factors contributing to this variability, we designed a large-scale mouse experiment in one animal facility that incorporated standardized phenotyping, microbiome analysis and metadata collection.

We administered 1.5% DSS through the drinking water for 7 days to 579 identically housed, SPF wild type C57BL/6N mice (297 male, 282 female; median 11 weeks; SD 12 days), from 14 distinct parental lineages (i.e. founder matings derived from a single breeding pair). We observed extensive variability in experimental endpoints in these genetically identical mice at 10 days post DSS treatment, with histological assessment of intestinal pathology of the mid and distal colon ranging from severe inflammation, characterised by extensive crypt loss, leukocytic infiltration and edema, to no visible inflammation (Fig. 1a, 1b). Substantial variation was also observed in the weight change of mice treated with DSS, with the response ranging from a 12% weight gain to a 30% weight loss across the cohort (Fig. 1c). Principal component and regression analysis considering histology scores and weight loss independently demonstrated limited correlation (R^2^ = 0.27; Fisher Exact Test) suggesting highly variable, multifactorial determinants in disease outcome in this experimental model (Fig. 1d). This also indicates that weight loss and intestinal pathology outcomes, common experimental endpoint measures in the DSS model, may be partially-independent responses to DSS.

**Figure 1:**
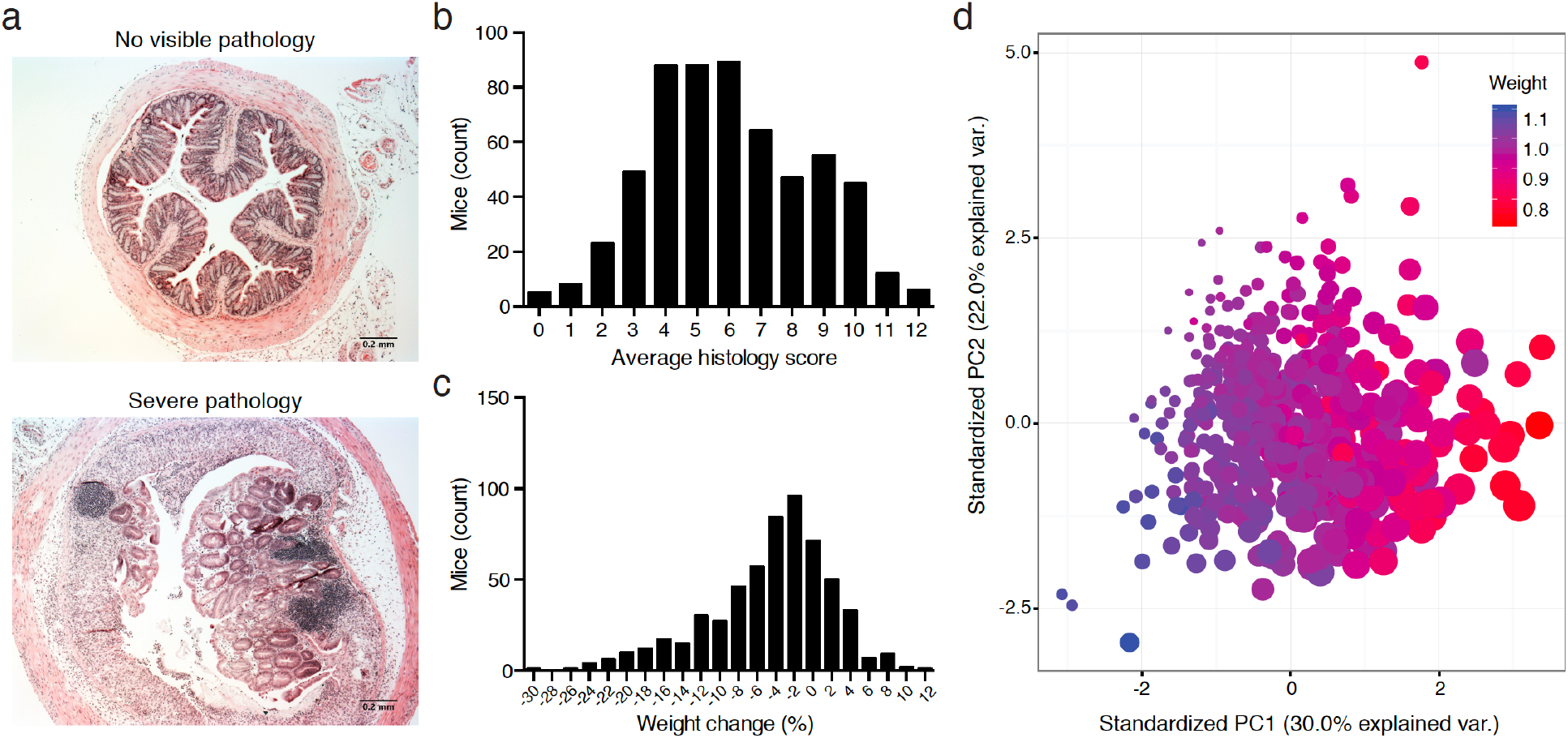
Disease severity is highly variable in the DSS mouse model. (a) Representative images of extremes of mid-colon inflammatory response of wild-type C57/BL6 mice after treatment with DSS showing a score of 0 and a score of 12. Distribution of (b) average histology scores and (c) weight change across 579 (297 male, 282 female; median 11 weeks; SD 12 days) wild-type C57BL/6N mice exposed to DSS treatment. (d) Principal component analysis considering the relationship between low weight loss (blue) and high weight loss (red) and increased inflammation (large) and minimal inflammation (small) demonstrates limited association across samples.

### Gut microbiota is major driver of DSS variability in genetically-identical mice

To identify the primary factors determining disease outcome, we established a random forest classifier model incorporating mouse sex, age, lineage, parents, mouse room and experimental date to assess the relative importance of each of these factors. The strongest predictive factors for weight loss and histology score, both together and independently in these genetically identical C57BL/6N mice, maintained under standardized SPF conditions within the same facility, was the parental lineage, with enrichment analysis confirming these associations (p < 0.01; Fisher Exact Test; Supplementary Fig. 1).

Since all the mice were derived from the same genetic founder stock the most likely mediator of their DSS phenotype was an extrinsic lineage specific factor. However, to exclude the role of host genetics or epigenetic differences between colonies, and to determine if the microbiota alone could be responsible for the observed phenotypic variation, faecal samples were collected from SPF mice (without DSS treatment) from parental lineages that represented extremes of disease outcome and used to colonise C57BL/6N germ-free mice. These colonised recipient mice were then subjected to DSS treatment with daily weight measurements with colon sections collected and analysed histologically 10 days after the initiation of DSS treatment. The recipient mice exhibited both weight loss and pathology scores that were not statistically different from those observed in the respective donor mouse lineages but differed significantly between disease-associated and non-associated lineages (p < 0.05; Fisher Exact Test) (Fig. 2a). From these results it is clear that, independent of host genetics or epigenetic lineage specific factors, the microbiota is sufficient to produce the observed variation in phenotypic response to DSS administration.

**Figure 2:**
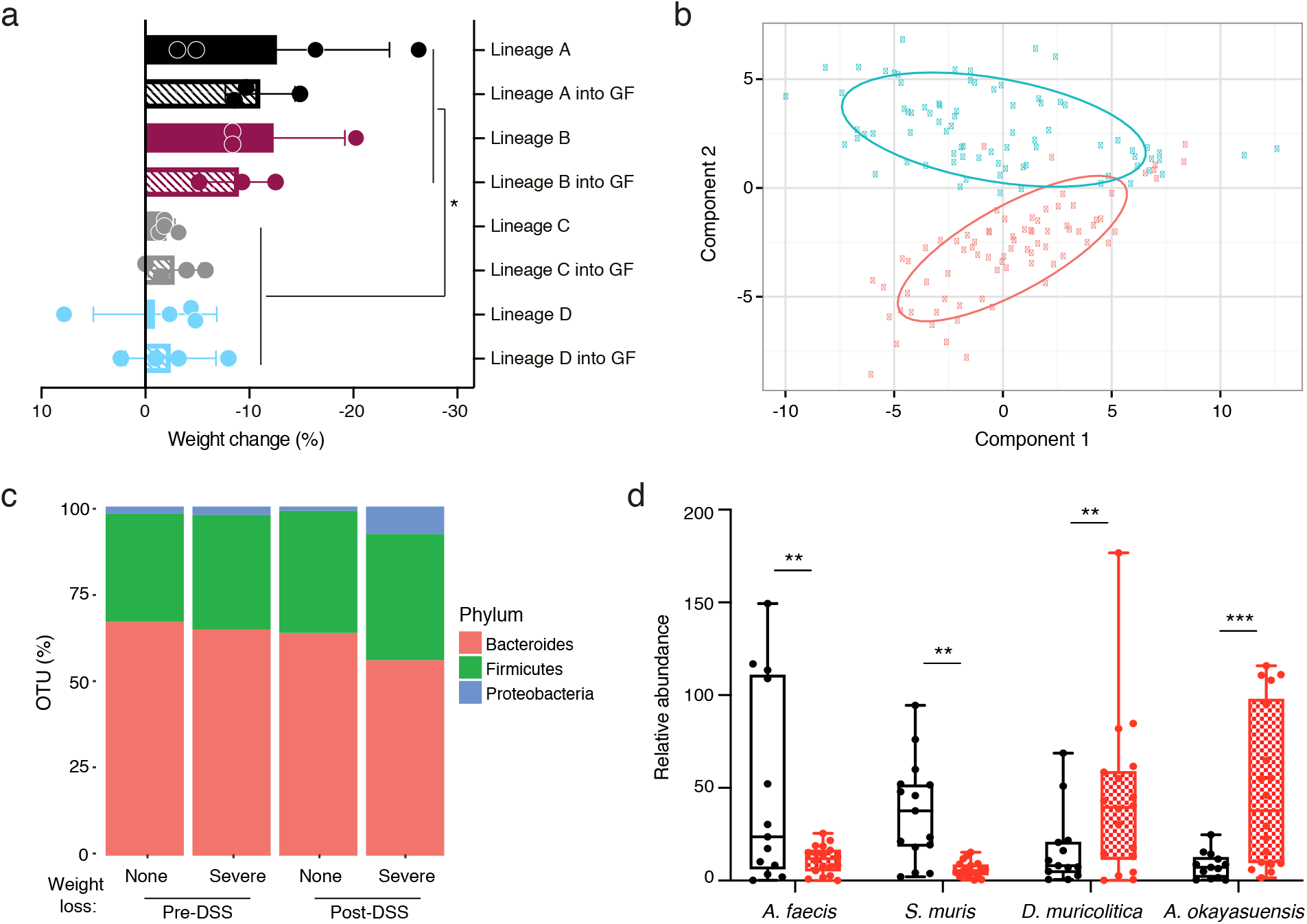
Discovery of candidate bacterial taxa associated with DSS disease variability. (a) Comparison of weight changed observed in conventional SPF mice (Lineage A, B, C and D; solid) and germ-free mice colonized by oral gavage (Lineage A, B, C and D into GF; hatched) after 10 day DSS challenge. (b) Principal component analysis of metagenomics profiling identifies a clear change between pre-DSS samples (blue) and post-DSS samples (red) regardless of lineage (c) Distribution of Bacteroides (red), Firmicutes (green) and Proteobacteria (blue) in samples that exhibited asymptomatic or severe weight loss and inflammation. (d) Relative abundance of *A faecis, A okayasuensis, D muricolitica and S. muris* (* p < 0.05; ** p < 0.01; *** p < 0.001; Mann Whitney test).

### Discovery of candidate bacteria that drive disease variability

Having established the importance of the microbiota in mediating phenotypic outcomes to DSS in genetically identical mice, we next sought to understand the degree of variability in mouse microbiota across a single facility and define the changes in the microbiota that occur with DSS treatment. Faecal samples were collected prior to DSS treatment and 10 days after DSS exposure, and subjected to taxonomic profiling with 16S rRNA gene sequencing to determine the microbiota community composition. Consistent with previous reports of gastrointestinal inflammation, this analyses demonstrated a clear separation of microbiome community between the pre- and post-DSS samples (Fig. 2b) and a significant reduction in bacterial diversity (Shannon diversity index; p < 0.05; Supplementary Figure 2)^14^. Expansion of Proteobacteria is often associated with gut microbiome dysbiosis, however, while an overall increase in Proteobacteria was observed, we did not observe a substantially greater increase in Proteobacteria levels in lineages that exhibited severe pathology or median weight loss over 5% compared to asymptomatic lineages (Fig 2c). This suggests other gut bacteria are responsible for the observed variation in DSS resistance and susceptibility.

Given the transmissibility of disease phenotype we observed in the germ-free mouse experiments, we reasoned the observed phenotypic variation was primarily dependent on the baseline microbiota composition, prior to DSS treatment. To understand the composition of the microbiota that was associated with increased risk of disease outcome we applied linear discriminant analysis to the gut microbiota composition prior to and after DSS exposure^15^. This unsupervised analysis leveraged the variability between mouse lineage-specific bacteria to identify the bacterial taxa that most closely predict DSS response independent of starting community. Using this approach, we identified two statistically different bacterial taxa associated with no disease and five statistically different bacterial taxa associated with severe disease. Notably, no equivalent predictive bacterial taxa were identified within the microbiota of mice post-DSS exposure. Our results suggest that the presence of specific gut bacterial taxa prior to DSS exposure influences disease outcome, possibly by either priming severe disease (pathobiont) or protecting from severe disease (commensal).

### Culture and isolation of candidate bacterial species to enable functional validation

While we show clear associations between specific bacterial taxa and DSS-mediated disease outcome using 16S rRNA gene sequencing, accurate taxonomic identification to strain level and experimental validation requires the isolation and genome sequencing of pure clonal cultures of candidate bacterial strains. To target and purify the candidate bacterial strains, we next performed broad culturing of ∼2000 bacterial colonies from mouse faecal samples. The resulting isolates were identified by capillary 16S rRNA gene sequencing, and included four of the previously identified bacterial taxa, 2 “disease-associated” and 2 “health-associated” (Fig. 2d). These isolates were subjected to whole genome sequencing and genomic analysis to aid in precise taxonomic and phylogenetic assignment (Supplementary Fig. 3). In addition, we performed extensive phenotypic characterization and comparative genome analyses of these 4 species (see Methods) as a basis to provisionally name the health-associated bacteria *Anaerostipes faecis* and *Sangeribacter muris* and the disease-associated bacteria *Duncaniella muricolitica* and *Alistipes okayasuensis*. Of note, the latter 3 of these species are mouse-specific.

We next repeated the DSS experimental model in germ-free mice or germ-free mice stably mono-colonised for four weeks with one of these four candidate bacteria (Supplementary Fig. 4a). Control germ-free mice treated with DSS did not lose weight (Fig. 3a) nor experience mortality (Fig. 3b) over the course of the experiment, suggesting that the presence of gut bacteria is important to trigger overt disease in our facility (Fig. 3). Comparable experimental outcomes were observed in DSS-treated germ-free mice mono-colonised with either of the health-associated bacteria, which did not lose weight or develop intestinal pathology (Fig. 3), nor experience significant mortality (Fig. 3b). In contrast, when we DSS treated mice mono-colonised with either of the disease-associated bacteria we observed more rapid and greater weight loss (Fig. 3a) and significantly reduced survival (Fig. 3b). Therefore, using a Koch’s Postulates approach^16,17^, we identified and validated *D. muricolitica* and *A. okayasuensis* as mouse pathobionts that can promote severe disease not observed in mice mono-colonized with the commensal bacteria *S. muris* and *A. faecis*.

**Figure 3:**
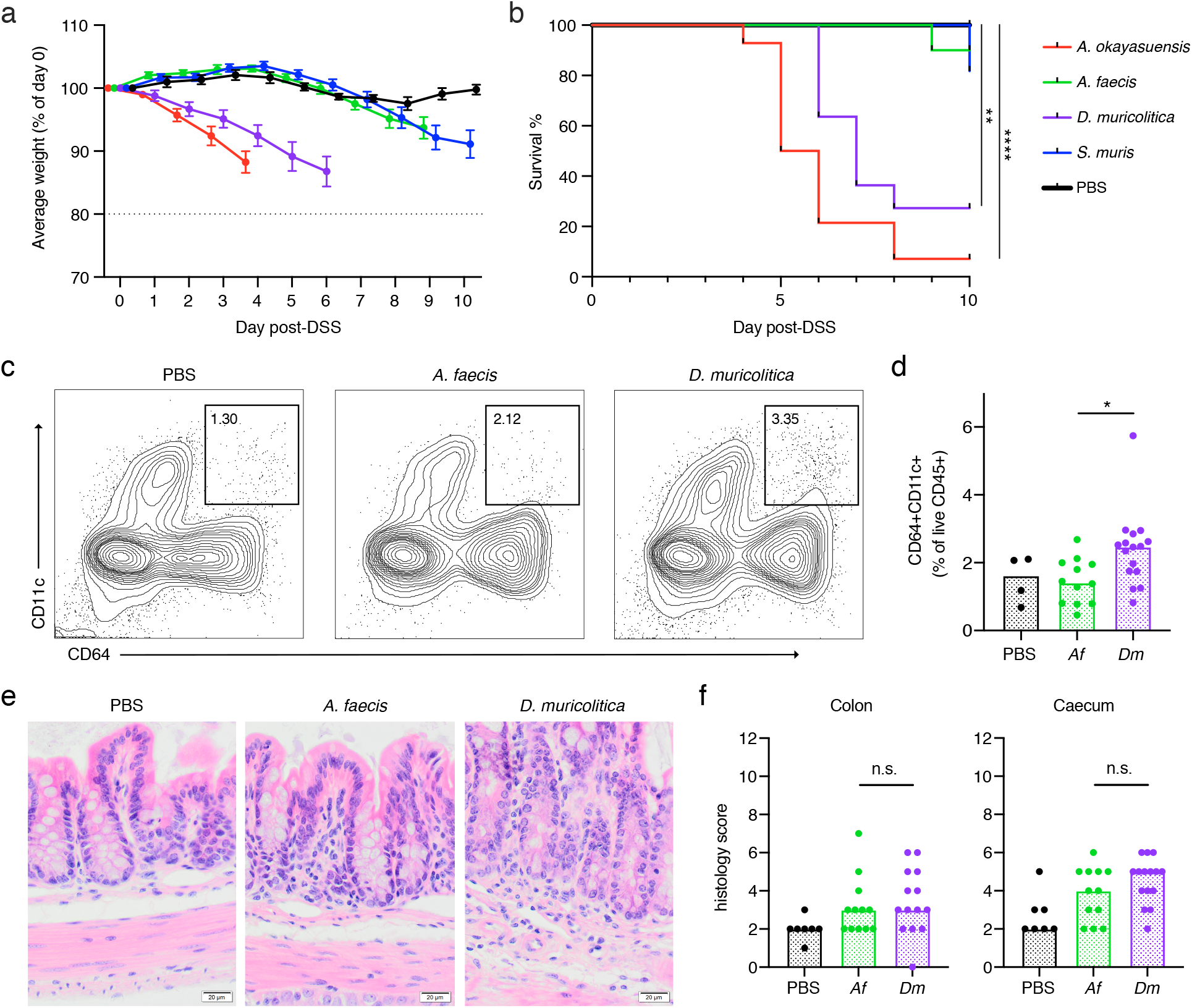
Phenotypic analysis of mice mono-colonised with candidate bacterial strains. (a) Average weight loss and (b) survival in gnotobiotic mice pre-colonised with either *A. okayasuensis* (red), *A faecis* (green), *D. muricolitica* (purple) or *S. muris* (blue) and germ-free controls (PBS; black) in response to oral DSS challenge. (c) Flow cytometric analysis of monocytes/macrophages (B220/CD3e/TCRg-MHC2+CD64+CD11c+) from the large intestines of control germ-free mice (PBS) or mice monocolonised with *A. faecis* or *D. muricolitica* and challenged with oral DSS for 7 days. (d) Lamina propria monocytes/macrophages as a percentage of viable CD45+ cells. (e) Representative colon images (400X) and (f) histological scores from the H & E staining of mid-colon and caecum of control germ-free mice (PBS) or mice mono-colonised with *A. faecis* or *D. muricolitica* and challenged with oral DSS for 7 days. Average weight loss curves were censored after the first mouse in each group reached 80% of baseline weight. For (b) all survival curves were compared to PBS control by log-rank (Mantel-Cox) test. For (d) and (f) the two colonisation conditions were compared to each other by Mann Whitney test. (* p < 0.05; ** p < 0.01, ****p<0.0001)

To understand the inflammatory immune responses in the intestines potentially caused by disease-associated versus health-associated species, we next examined immune cell populations and cytokine production. Flow cytometric analysis of intestinal lamina propria leukocytes isolated from germ-free mice mono-colonised for 4 weeks with either commensal *A. faecis* or pathobiont *D. muricolitica* revealed that colonisation with either strain did not significantly impact the composition of intestinal leukocyte subsets (Supplementary Fig. 4b, c). However, the perturbation and epithelial disruption induced by DSS led to significantly increased inflammatory CD64+CD11c+ monocyte/macrophages in the lamina propria compartment of the large intestine in *D. muricolitica* mono-colonised mice, but not in *A. faecis* mono-colonised mice (Fig. 3c, d). These monocytes have been shown to play a pathogenic role in the intestines in the context of *Helicobacter*-induced inflammation^18^. This suggests that, although not pathogenic or immunogenic at homeostasis, *D. muricolitica* drives increased inflammatory responses in the wake of epithelial damage whereas health-associated *A. faecis* does not. Importantly, while histological samples taken after 7 days of DSS treatment from colons and caecums of *D. muricolitica* mono-colonised mice occasionally exhibited features of severe inflammation, this was not statistically different from that of *A. faecis* mono-colonised mice (Fig. 3e, f) and did not consistently correspond with the weight loss observed. This again suggests that weight loss in the DSS model may not always correlate with intestinal inflammation and pathology.

### Global distribution and prevalence of *D. muricolitica* and *A. okayasuensis* in mouse facilities

As *D. muricolitica* and *A. okayasuensis* affected the outcome of DSS colitis in our mouse facility, we next explored the presence and distribution of these mouse pathobionts in other mouse facilities around the world. To do so, we first generated a global representation of the laboratory mouse intestinal microbiomes by curating 582 publicly available shotgun metagenome samples from “control”, SPF mice. We combined this dataset with shotgun metagenomes from faeces of 40 mice at the Wellcome Sanger Institute to yield a global mouse gut microbiome dataset that covers 31 institutes across 12 countries. We then determined the prevalence and abundance of the health-associated bacteria *S. muris* and *A. faecis* and disease-associated bacteria *D. muricolitica* and *A. okayasuensis* in this mouse microbiome dataset.

*D. muricolitica* and *A. okayasuensis* were both dominant members of the mouse microbiome, prevalent in 54.5% and 47.9% of samples with a mean abundance of 0.52% and 0.48% reads per sample respectively. Importantly, *D. muricolitica* and *A. okayasuensis* were each detected in 80.6% (25/31) of the institutes included in the analyses (Figure 4), and there were only 3 institutes that did not contain either of the pathobionts. For comparison, of the health-associated species, *S. muris* was the most dominant species of the mouse gut microbiota with a prevalence of 96.5% and a mean abundance of 6.2% reads, while *A. faecis* was a relatively minor species, with a prevalence of 5.9% and a mean abundance of 0.22% reads. *S. muris* was present in all (31/31) of the institutes included in the analyses, whereas *A. faecis* was detected in 25.8% (8/31) of the institutes. The abundance of each species also differed between institutes (Figure 4). Thus, we show that mouse-derived bacteria cultured from our insitute are common mouse symbionts and, importantly, the pathobionts *D. muricolitica* and *A. okayasuensis* are common, but not ubiquitous, in animal facilities around the world.

**Figure 4:**
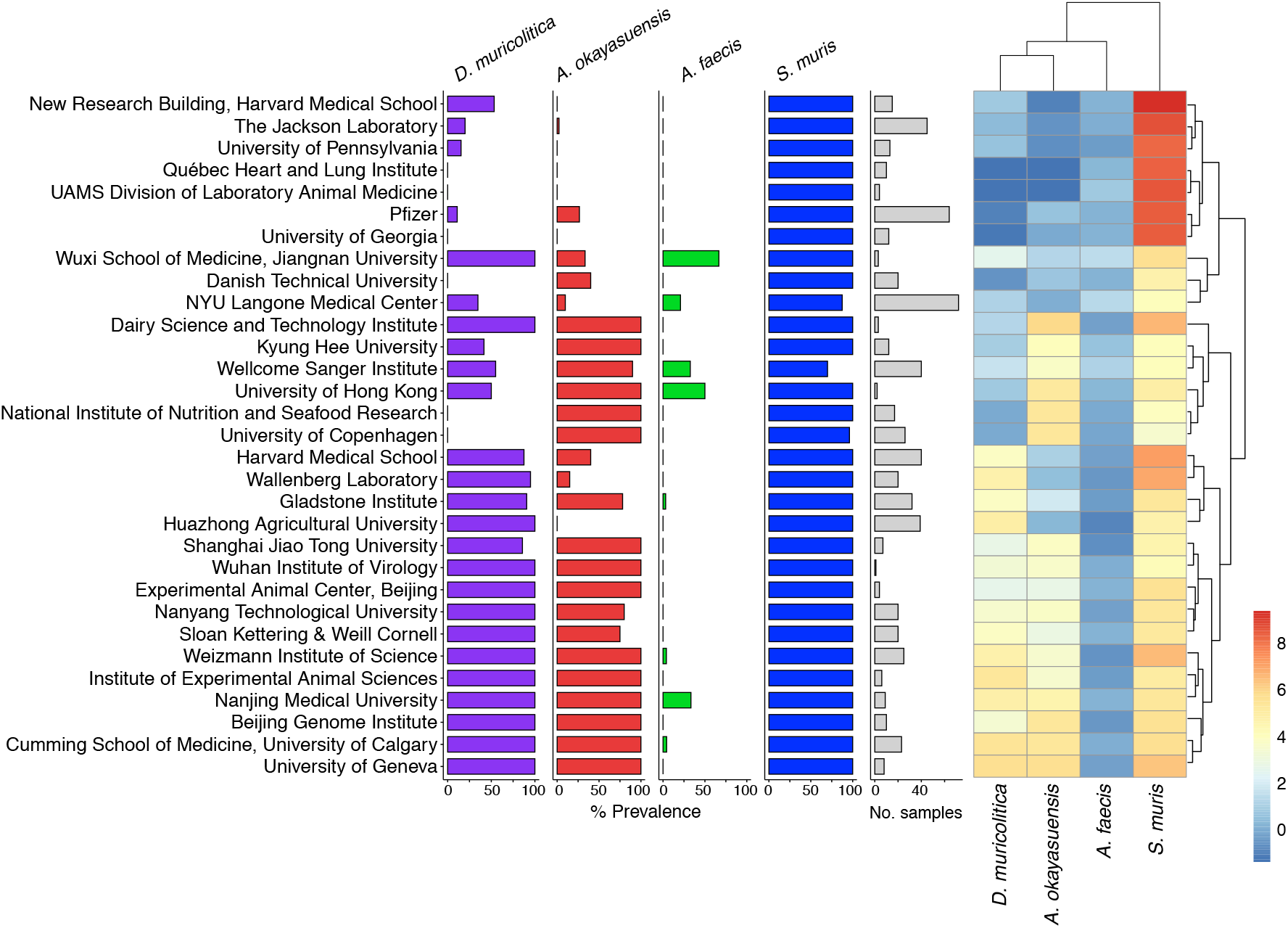
Global epidemiology and intra-institutional variation of disease- and health-associated species. Prevalence and abundance of disease- and health-associated species in SPF mice across international mouse facilities. Left to right: intra-institute prevalence of *D. muricolitica, A. okayasuensis, A. faecis* and *S. muris*; bar plot indicating number of samples per institute; heatmap of mean abundance for each species. A threshold of ≥0.01% metagenomic reads is used to define prevalence of a species. For mean abundance, count zeros are removed using Bayesian multiplicative replacement and data scaled using a centered log-ratio transformation.

## DISCUSSION

Despite experimental variability of the DSS mouse model, which limits the interpretation and translation of data, the model is increasingly used in basic research (>6000 DSS model publications on PubMed) and by the pharmaceutical and biotechnology industries. Co-housing, heterozygote mating and other experimental procedures have been used in an attempt to normalize the microbiota within experiments and ensure direct measures of the effects of mouse genetic background on disease phenotype rather than indirect and unknown microbiota-associated effects^19^. In our approach, we leveraged this variability in DSS outcome, combined with microbiome analysis, machine learning, anaerobic culturing and germ-free mouse validation experiments to gain relevant biological insight into microbiota determinants of DSS disease. Importantly, we identified causative bacterial strains *D. muricolitica* and *A. okayasuensis* and fulfilled Koch’s postulates.

Interestingly, *D. muricolitica* and *A. okayasuensis* are phylogenetically distinct from each other, and neither carry known virulence factors. As such, it remains unclear how they contribute to DSS disease, raising the need for further experimental study of host interactions and the biology of these important mouse specific bacteria. We have also noted that weight loss and intestinal pathology do not always correlate, so they should not necessarily be used as surrogates for each other.

We propose that researchers should monitor for *D. muricolitica* and *A. okayasuensis* in their mouse facilities, as is undertaken for other immunophenotyped-affecting microbes such as segmented filamentous bacteria^20^. In some cases, pre-colonisation of animals with these bacteria if they are not present in the facility may yield better experimental control and increase confidence in this important and widely used mouse model of human disease. Overall, our work suggests application of metagenomics techniques to report the microbiota composition in addition to the genetic and disease phenotypes being described should represent a standard minimum requirement for the DSS mouse model. We have deposited the *D. muricolitis* and *A. okayasuensis* strains and genomes in public repositories to support this work.

## Methods

### Mouse models

Mice were maintained under specific pathogen-free conditions at a Home Office-approved facility with all procedures carried out in accordance with the United Kingdom Animals (Scientific Procedures) Act of 1986. DSS was administered through the drinking water at a concentration of 2% to C57BL6/6N mice aged between 7 and 16 weeks (297 male, 282 female; median 11 weeks; SD 12 days). Faecal transplant and single colonised gnotobiotic lines were generated by weekly oral gavage over a 3-week period. Mice were maintained within individually ventilated cages.

### DSS colitis challenge

Colitis was induced by adding 1.5% (w/v) DSS (Affymetrix, Inc.) to drinking water for 7 days, followed by 3 days with regular drinking water, in animals aged between 5 and 18 weeks (mean age 9 weeks). Mice were weighed every day and culled if weight loss reached 20% of starting weight. Average weight loss curves were censored for each group when the first member of the group reached this end-point to avoid misleading calculations of the average using the remaining members.

### Histological assessment of intestinal inflammation

DSS-treated SPF mice were sacrificed at day 10 by cervical dislocation, and samples from mid and distal colon taken. Tissue sections were fixed in buffered 10% formalin; paraffin-embedded; cut; and stained with haematoxylin and eosin. Colon histopathology was blind-graded semi-quantitatively on a scale from zero to three, for four criteria: (1) degree of epithelial hyperplasia/damage and goblet cell depletion, (2) leukocyte infiltration in lamina propria, (3) area of tissue affected, and (4) presence of markers of severe inflammation, including crypt abscesses, submucosal inflammation, and oedema. Scores for individual criteria were added for an overall inflammation score of between zero and twelve for each sample. Scores from mid and distal colon were then averaged to obtain inflammation scores for each mouse colon.

For monocolonised germ-free mice, animals were euthanized 7 days after DSS administration, and samples were taken from the caecum and mid colon. Tissue sections were fixed in methacarn, paraffin-embedded, sectioned, and stained with haematoxylin and eosin. Colon and caecum histopathology was graded on a semi-quantitative scale by a pathologist blinded to the groups. The criteria scored were 1) degree of epithelial hyperplasia/damage and goblet cell depletion, 2) severity of leukocyte infiltration in lamina propria 3) extent of the inflammation, and 4) presence of markers of severe inflammation, including crypt abscesses, submucosal inflammation and oedema. Each criterion was scored on a scale of zero to three based on previously described thresholds^21^ and the sum of the individual scores was recorded as an indication of the overall inflammation.

### Bacterial Culturing

Faecal samples and bacterial isolates were cultured as described previously for anaerobic gastrointestinal bacteria under anaerobic conditions (10% H, 10% CO2, 80% N) in a Whitley DG250 workstation at 37 °C^8^. Faecal samples were homogenized in reduced PBS (0.1 g per ml PBS), serially diluted and plated directly onto YCFA agar supplemented with 0.002 g ml−1 each of glucose, maltose and cellobiose. Colonies were picked, restreaked to purity and identified using 16S rRNA gene sequencing.

### Microbiota profiling and analysis

Microbiota assessment was performed using the FastDNA Spin Kit for Soil (MP Biomedicals) on 300mg of faecal sample from each mouse. The V1-V2 region of the 16S rRNA genes was amplified with Q5 High-Fidelity Polymerase Kit (New England Biolabs). (F’:AATGATACGGCGACCACCGAGATCTACAC-TATGGTAATT-CC-AGMGTTYGATYMTGGCTCAG; R’:CAAGCAGAAGACGGCATACGAGATACGAGACTGATTAGTCAGTCAGAAGCTGCCTCCCGTAGGAG). Each PCR amplification was performed as four independent amplifications and pooled in equimolar amounts for 150bp paired end sequencing with the Illumina MiSeq platform. Resulting 16S rRNA amplicon sequences were analysed with mothur MiSeq SOP v.1.42.3 using SILVA v.132^22^.

The 16S rRNA gene alignments were used to determine a maximum likelihood phylogeny using FastTree v.2.1.10^23^. Phylogenetic trees were visualized and edited using iTOL^24^. Principal component analyses were performed in R and Linear regression analysis was performed using python scikit-learn 0.20.4. Alpha diversity was assessed using Vegan 2.5.6.

### Intestinal immune cell isolation

Large intestines were excised and separated from mesentery and fat. Transluminal sections were taken from the caecum and distal colon for histology. The intestines were opened longitudinally, cleared of faeces, and then washed three times in cold PBS. To isolate the intraepithelial immune populations, the large intestines were cut into 1cm pieces, washed in cold PBS, and then incubated in 10 mL of PBS + 1 mM dithiothreitol for 10 minutes. Tissues were manually disrupted via shaking and then strained, collecting the supernatant. Tissues were then incubated in 10 ml of PBS + 30 mM EDTA + 10 mL HEPES at 37°C at 200 rpm, before being shaken and strained. The supernatants from these two steps were pooled, filtered at 70 µm and the filtrate fractionated using a discontinuous Percoll gradient (80%/40%). Epithelial cells were isolated from the surface of the Percoll, and the intraepithelial immune cells isolated from the interface.

To access the lamina propria compartment, the tissues were manually chopped and digested in HBSS + 25 mM HEPES + 1 mM sodium pyruvate containing 0.05 mg/mL Collagenase VIII (Sigma) and 50 µg/ml DNase I (Sigma) for 1 hour at 37°C at 80 rpm. Samples were mechanically disrupted and filtered at 70 µm. The filtrate was fractionated using a discontinuous Percoll gradient (80%/40%). Lamina propria immune cells were isolated from the interface.

### Flow cytometry

Intestinal and mesenteric lymph node immune cell populations were characterised by flow cytometry. For intracellular cytokine staining, cells were incubated in T cell media (RPMI, 10% FBS, 1 mM sodium pyruvate, 2mM GLUTamax, 1X non-essential amino acids, 0.1 mM 2-β-mercaptoethanol, 10 mM HEPES, 1%Penicillin/Streptomycin) + 1X Cell Stimulation Cocktail with Protein Inhibitors (eBioscience) for 3 hours at 37°C before staining. Non-viable cells were stained using Live/Dead Fixable Aqua. Cells were permeabilised with CytoFix/CytoPerm (BD) followed by intracellular staining in PermBuffer (eBioscience).

All antibodies were purchased from eBioscience unless otherwise indicated. Antibodies used were CD45-SB600, TCRb-APC-Cy7, MHCII-FITC, CD4-PE-Cy7, CD8a-AF700, CD8b-PE, IFNg-PerCP-Cy5.5, TNFa-APC, IL-17A-bv421 (BioLegend), CD3e-FITC, TCRg-FITC, B220-FITC, MHCII-APC-e780, CD11b-PerCP-Cy5.5, CD11c-AF700, CD64-APC (BioLegend), SiglecF-PE (BD), Ly6G-Pe-Cy7 (BD) and F4/80-e450. Sample data were acquired with an Attune NxT flow cytometer coupled with an Attune CytKick Max autosampler. Data were analysed using FlowJo v10.

### Curation of public metagenome samples

To assess the abundance of the health- and disease-associated bacteria in mice from different institutes, we curated publicly available faecal shotgun metagenomes for healthy, “control”, SPF mice from NCBI and the ENA. These samples included mice from different genetic backgrounds and strains, but samples were excluded if mice were younger than 3 weeks, had received antimicrobial treatment or dietary intervention, or had a gene knockout. In total 582 faecal shotgun metagenomes were curated.

### Sample collection and Shotgun metagenomic sequencing

Faecal samples were collected directly from mice, and immediately stored at -80°C until DNA extraction. DNA was extracted from faecal samples using FastDNA Spin Kit for Soil (MPBio) and stored at -20 °C until metagenomic sequencing. DNA samples were quantified using a Qubit 4 Fluorometer (Thermo Fisher), and samples with >100 ng DNA material proceeded to paired-end (2 × 150 bp) metagenomics sequencing on the HiSeq 4000 platform.

### Assessment of Microbial Abundance Across Facilities

Metagenomes were quality controlled using KneadData v0.7.3 with default settings. Host reads were removed from samples using the GRCm39 reference genome and Bowtie2 v2.3.5. In addition, reads were aligned to the Phi X 174 genome and removed. Taxonomic classification of metagenomics reads was performed using Kraken2 v2.0.8 against genomes of the Mouse Microbial Genome Collection. The relative abundances of species were corrected using Bracken v2.5.2. Data analysis was performed in R v4.0.2. Species abundant at ≥0.01% reads were considered prevalent in a sample. To quantify and visualise intra-institute abundance, Bayesian multiplicative replacement of count zeros was performed using the zCompositions v1.3.4 package, and data transformed using centre-log ratio normalisation. Mean abundance was then visualised using the pheatmap v1.0.12 package.

### Novel species characterisations

While metagenome assembled genomes are publicly available for the species described in this paper, and they are represented by genomic species clusters in the Genome Taxonomy Database (A14=Anaerostipes sp000508985, A43=CAG-485 sp002362485, A60=Duncaniella sp001689575, A61=Alistipes sp002362235), these species have not yet been characterised nor have these names been validly published according to the International Code of Nomenclature of Prokaryotes. As these species are uncharacterised they are referred to as novel in this manuscript. We combined single carbon source utilisation data, generated using the Biolog platform (data in Supplementary Table 1), with genome annotation by eggNOG-mapper v2 and InterProScan v5 to predict encoded functionality in order to characterise these novel species. We describe these species and propose new names below.

#### Description of Alistipes okayasuensis sp. nov

*Alistipes okayasuensis* (o’ka.ya.su.en.sis. N.L. masc./fem. adj. *okayasuensis*, named after Isao Okayasu, who first described dextran sulphate sodium colitis as a model for ulcerative colitis in mice. Phylogenomic analyses place strain A61^T^ in the *Alistipes* cluster. The dDDH value between *Alistipes onderdonkii* and *Alistipes putredinis* is 24.4%, while the dDDH values between strain A61^T^, *Alistipes onderdonkii* and *Alistipes putredinis* are 25.2% and 21.4% respectively. The G+C content difference between the genome of strain A61^T^ and *Alistipes onderdonkii*, its closest phylogenomic neighbour, is 1.39%, while their 16S rRNA gene sequence identity is 96.03%. These analyses strongly indicate that *Alistipes okayasuensis* is a separate species within the genus *Alistipes*. This strain is a strict anaerobe. Carbon source utilisation analysis was combined with reconstruction of KEGG metabolic pathways (55.58% of 2022 predicted ORFs could be annotated) to functionally characterise the strain. The strain encodes 47 genes predicted to be carbohydrate-active enzymes (CAZymes), including 29 glycoside hydrolases: seven beta-glucosidases (GH3, EC 3.2.1.21); five beta-galactosidases (EC 3.2.1.23); five endohydrolytic alpha-glucosidases (GH13) including four alpha-amylases (EC 3.2.1.1); three exohydrolytic alpha-glucosidases (GH31, EC 3.2.1.20); three cellulases (EC 3.2.1.4); three mannan endo-1,4-beta-mannosidase (EC 3.2.1.78); and an alpha-L-fucosidase (EC 3.2.1.51). Despite this predicted hydrolytic potential, the strain shows limited capacity to use disaccharides and oligosaccharides as sole carbon sources. While it can utilise alpha-D-lactose, sucrose and maltotriose, it cannot grow on D-cellobiose, dextrin, cyclodextrin, gentiobiose, lactulose, maltose, D-melezitose, D-melibiose, palatinose, D-raffinose, stachyose, D-trehalose, fucose or turanose. The strain has enzymes for metabolism of L-histidine to L-glutamate via 4-Imidazolone-5-phsophate and is able to use urocanate as a sole carbon source (EC 4.3.1.3, EC 4.2.1.49, EC 3.5.2.7, EC 2.1.2.5). In keeping with some other members of the *Alistipes* genus, this strain encodes a putative tryptophanase (EC 4.1.99.1) that allows it to hydrolyse tryptophan to indole. The strain encodes a butyrate phosphotransferase (EC 2.3.1.19) and a butyrate kinase (2.7.2.7), indicating potential to metabolise butyrate. The genome is 2,329,524 bp with a G+C content of 56.45 mol%. We propose the name *Alistipes okayasuensis* for this novel species, in reference to the origins of the DSS colitis model and reflecting this species’ implication in pathogenesis.

#### Description of Anaerostipes faecis sp. nov

*Anaerostipes faecis* (fae’cis. L. gen. fem. n. *faecis*, of faeces, referring to faecal origin). The closest phylogenomic neighbour of strain A14^T^ is *Anaerostipes caccae* with a dDDH value of 48.40%. The pairwise 16S rRNA gene sequence identity of A14^T^ and *A. caccae* DSM 14662^T^ is 98.16%. These analyses indicate that A14^T^ is a separate species within the genus *Anaerostipes*. The genome is 3,200,246 bp with a G+C content of 43.77 mol%. We propose the name *Anaerostipes faecis* for this new species.

The strain is strictly anaerobic. Reconstruction of KEGG metabolic pathways (55.4% of 3222 predicted ORFs could be annotated) indicates that the strain produces butyrate via a butyrate phosphotransferase/butyrate kinase operon (EC 2.7.2.7, EC 2.3.1.91), in keeping with the genus type species *A. caccae*. Likewise, the strain encodes an arginine dihydrolase (EC 3.5.3.6). In contrast to the genus description however, this strain does not encode an alpha-galactosidase (EC 3.2.1.22) nor a phosphoamidase (EC 3.9.1.1).

The strain shows limited capacity to utilise monosaccharides as sole carbon sources: it reliably metabolises 3-methyl-D-glucose, but did not utilise any other monosaccharides include D-fructose, D-galactose, D-mannose or D-glucose. Notably, the strain encodes the complete Leloir pathway for galactose metabolism as well as multiple beta-galactosidases (EC 3.2.1.23), indicating that the strain could metabolise galactose in different contexts. The strain can utilise some disaccharides, metabolising sucrose via multiple oligo-1,6-glucosidases (EC 3.2.1.10), D-trehalose via three trehalases (EC 3.2.1.28, GH37, GH65) and turanose, but does not utilise alpha-D-lactose, lactulose, maltose, D-melibiose, gentiobiose, D-cellobiose or palatinose. The strain could not metabolise the trisccharides maltotriose, D-melezitose, D-raffinose, nor the tetrasaccharide stachyose – potentially due to the absence of an alpha-galactosidase. The strain could utilise beta-cyclodextrin, but not dextrin or alpha-cyclodextrin. The strain is capable of metabolising alpha-ketovaleric acid and D-malic acid, as well as certain amino acids and derivatives including L-threonine, L-asparagine, L-serine and the dipeptides glycyl-L-proline and L-alanyl-Lthreonine. In our study, it could not metabolise L-glutamine, L-glutamate, L-valine nor L-alanine.

#### Description of Duncaniella muricolitica sp. nov

*Duncaniella muricolitica* (mu.ri.co’li.ti.ca. L. gen. n. *muris*, of the mouse; Gr. n. *kolon*, colon; Gr. suff. *- tikos -ê -on*, suffix used with the sense of pertaining to; N.L. fem. adj. *muricolitica*, pertaining to the colon of the mouse). The closest phylogenetic neighbour according to pairwise 16S rRNA gene analyses is *Duncaniella dubosii* with a sequence identity of 94.04%. The 16S gene sequence identity values between *D. dubosii* H5^T^, *D. muris* DSM 103720^T^ and *D. freteri* TLL-A3^T^ are 95.14% and 93.00% respectively, indicating that A60^T^ is a separate species within the *Duncaniella* cluster. The dDDH value between A60^T^ and *D. dubosii* H5^T^ is 28.10%, confirming a separate species status.

Recent work benchmarking an average nucleotide identity (ANI) approach to define genus boundaries found that the genus inflection points of members of the order *Bacteroidales* had relatively low values for alignment fraction (AF) and ANI (AF:ANI: *Bacteroides fragilis*=0.215:71.23; *Porphyromonas asaccharolytica*=0.105:68.12; *Prevotella melaninogenica*=0.195:72.07). The corresponding values between A60^T^ and the genus type species *D. muris* are 0.243:80.85, providing further evidence that A60^T^ belongs to the genus *Duncaniella*.

The strain is an obligate anaerobe. Reconstruction of KEGG metabolic pathways (47.2% of 3461 predicted ORFs could be annotated) indicated that strain is enriched for enzymes with glycoside hydrolase activity, encoding 47 in total. It is able to utilise dextrin, beta-cyclodextrin and various oligosaccharides including D-melezitose, maltotriose, amygdalin, maltose, gentiobiose (EC: 3.2.1.21), turanose, and D-cellobiose (EC: 2.4.1.20; EC: 3.2.1.21). The strain metabolises D-raffinose via sucrose (EC: 3.2.1.22, EC: 3.2.1.20), instead of via stachyose or melibiose, neither of which it can metabolise. In our study, the strain also did not utilise lactose as a single carbon source, even though it encodes seven predicted beta-galactosidases (EC 3.2.1.23). The strain is enriched for enzymes with glycosyltransferase activity, encoding 36 in total. Among these are seven GT2 family enzymes including two dolichyl-phosphate beta-D-mannosyltransferases (EC: 2.4.1.82).

The strain can utilise a variety of monosaccharides as single carbon sources, including alpha-D-glucose, alpha-D-glucose 1-phosphate, 3-methyl-D-glucose, D-galactose, alpha-methyl-D-galactose, D-mannose, adonitol, arabinose, D-gluconate, D-arabitol, and D-sorbitol. It did not metabolise fucose, alpha-D-lactose, D-fructose, lactate, lactulose, D-trehalose or L-rhamnose. The strain has enzymes for metabolism of L-histidine to L-glutamate via 4-Imidazolone-5-phsophate and is able to use urocanate as a sole carbon source (EC 4.3.1.3, EC 4.2.1.49, EC 3.5.2.7, EC 2.1.2.5). The genome is 4,122,701 bp with a G+C content of 50.9%. Following functional and genomic characterisation of our isolate, we propose the name *Duncaniella muricolitica* for this species, in reference to its association with poor prognosis in the murine DSS model of colitis. The genome is 4,054,788 bp with a G+C content of 50.93 mol%. Following functional and genomic characterisation of our isolate, we propose the name *Duncaniella muricolitica* for this species, in reference to its association with poor prognosis in the murine DSS model of colitis.

#### Description of Sangeribacter gen. nov

*Sangeribacter* (San.ge.ri.bac’ter. N.L. gen. masc. n. *Sangeri*, of Sanger; N.L. masc. n. *bacter*, rod; N.L. masc. n. *Sangeribacter*, a rod-shaped bacterium named after Frederick Sanger (1918-2013) and the institute where this genus was first described). 16S rRNA gene sequence analyses place this strain in the *Muribaculaceae* family, and *Sangeribacter* possess all features of this family. The pairwise sequence identity of the 16S rRNA gene of A43^T^ with 16S genes from *Paramuribaculum intestinale* DSM 100749^T^, *Muribaculum intestinale* YL27^T^ and *Duncaniella muris* DSM 103720^T^ is 88.90%, 88.22% and 87.39% respectively. The dDDH value between A43^T^ and *Paramuribaculum intestinale* DSM 100749^T^ is 35.5% and the difference in G+C mol% is 6.37%. Together these analyses strongly indicate that *Sangeribacter* is a separate genus. The type species *Sangeribacter muris* is one of the most dominant species in the mouse gut microbiota, representing up to 60% of classified reads in some mouse faecal shotgun metagenome samples. It is also highly prevalent with 77% positive samples of 1,926 in total.

#### Description of Sangeribacter muris sp. nov

*Sangeribacter muris* (mu’ris. L. gen. masc./fem. n. *muris*, of the mouse, the species was first isolated from a mouse). The strain is a strict anaerobe. It encodes 66 CAZy encoding genes including 42 glycoside hydrolases. It has a wide repertoire of glycans and carbohydrates that it can metabolise, including dextrin (EC 3.2.1.3), alpha- and beta-cyclodextrin, stachyose (EC 3.2.1.22), D-raffinose (EC 3.2.1.26), maltotriose, D-melezitose, amygdalin (EC 3.2.1.22), D-cellobiose (EC 3.2.1.21, EC 2.4.1.20), gentiobiose (EC 3.2.1.21), D-melibiose, palatinose, D-trehalose (EC 2.7.1.201, EC 2.4.1.64, EC 3.1.3.12), turanose, sucrose (EC 2.7.1.211), D-glucosaminate, lactulose, alpha-D-lactose, alpha-D-glucose, alpha-D-glucose 1-phosphate, 3-methyl-D-glucose, D-glucosaminic acid, alpha- and beta-methyl-D-glucoside, D-fructose (EC 2.7.1.4), D-mannitol (EC 1.1.1.67), D-mannose, D-galactose, alpha- and beta-methyl-D-galactoside, D-galacturonic acid, L-sorbitol (EC 1.1.1.14), L-fucose (EC 3.2.1.51). This strain could not utilise adonitol, dulcitol or D-gluconate as single carbon sources.

The strain is capable of metabolising amino acids and their derivatives, such as L-phenylalanine, L-asparagine, L-glutamate, L-glutamine, L-alanine, and L-valine, as well as dipeptides including L-alanyl-L-threonine, L-alanyl-L-glutamine, L-alanyl-L-histidine, glycyl-L-aspartate, glycyl-L-glutamine and glycyl-L-proline. It could not metabolise glycyl-L-proline nor utilise L-serine, L-aspartate or L-threonine as single carbon sources. The strain encodes a tryptophanase (EC 4.1.99.1) indicating that it can hydrolyse tryptophan to indole. The strain cannot utilise propionate or hydroxybutyrate, indicating that it is not a consumer of short chain fatty acids. The genome is 3,532,505 bp with a G+C content of 46.70 mol%. We propose the name *Sangeribacter muris* for this new species, reflecting its high abundance and prevalence in mice and its initial isolation from a murine host.

## Acknowledgements

This work was supported by the Wellcome Trust [098051] and the Australian National Health and Medical Research Council [1091097 and 1141564 to SF]. We are grateful to the Wellcome Sanger Institute Pathogen informatics and Research Support Facility for supporting this research. VP is supported by a Sir Henry Dale Fellowship jointly funded by the Wellcome Trust and the Royal Society [206245/Z/17/Z]. BBJ is supported by a studentship from the Rosetrees Trust [A2194]. KJM is supported by a Wellcome Trust Investigator Award (102972/Z/13/Z).

## Funding for open access charge

Wellcome Sanger Institute.

## Author Contributions

SC, TL, SF, VP conceived the study and study design. KH, SC, GN, CB, BB and AS performed animal experiments. SF, BB, NK undertook genomic and computational analysis. SF, GN, MS and AA cultured bacteria, BB, AS and VP performed mouse cell isolations and profiling. All authors contributed to study design and preparing the manuscript.

## Author Information

TDL is a founder and CSO of Microbiotica. The other authors declare no competing financial interests.

## Supplementary Figures

**Supplementary Figure 1:**
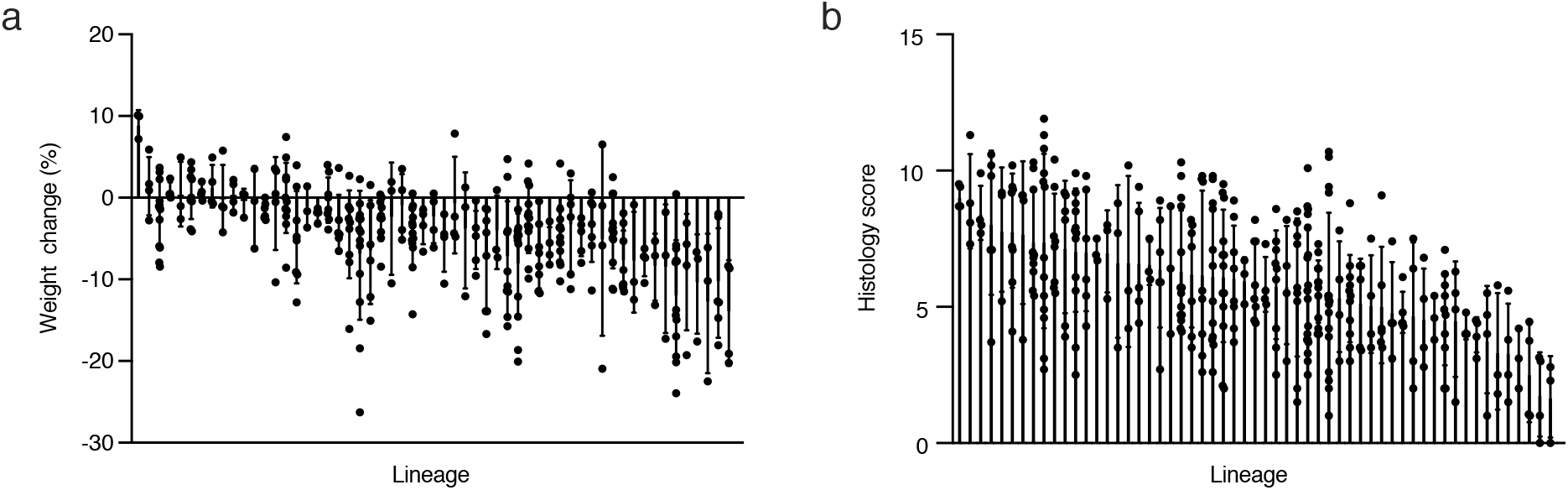
Lineage specific variation across C57BL/6 mice. Variation in (a) weight change and (b) average histology score across wild-type mouse lineages. Data presented as mean and standard deviation for each group.

**Supplementary Figure 2:**
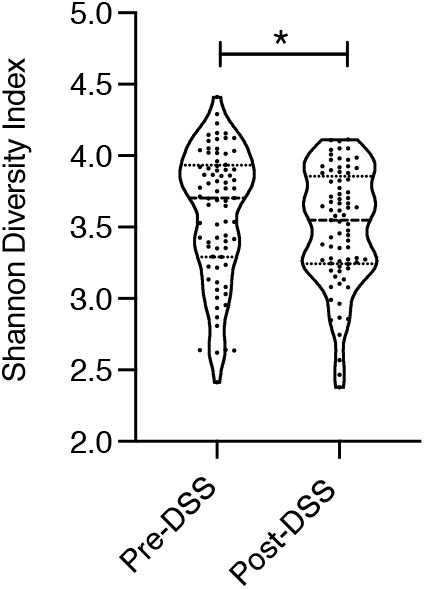
Shannon Diversity. Alpha diversity using Shannon-diversity index grouped by Pre-DSS and Post-DSS samples (* p < 0.05; paired T-test).

**Supplementary Figure 3:**
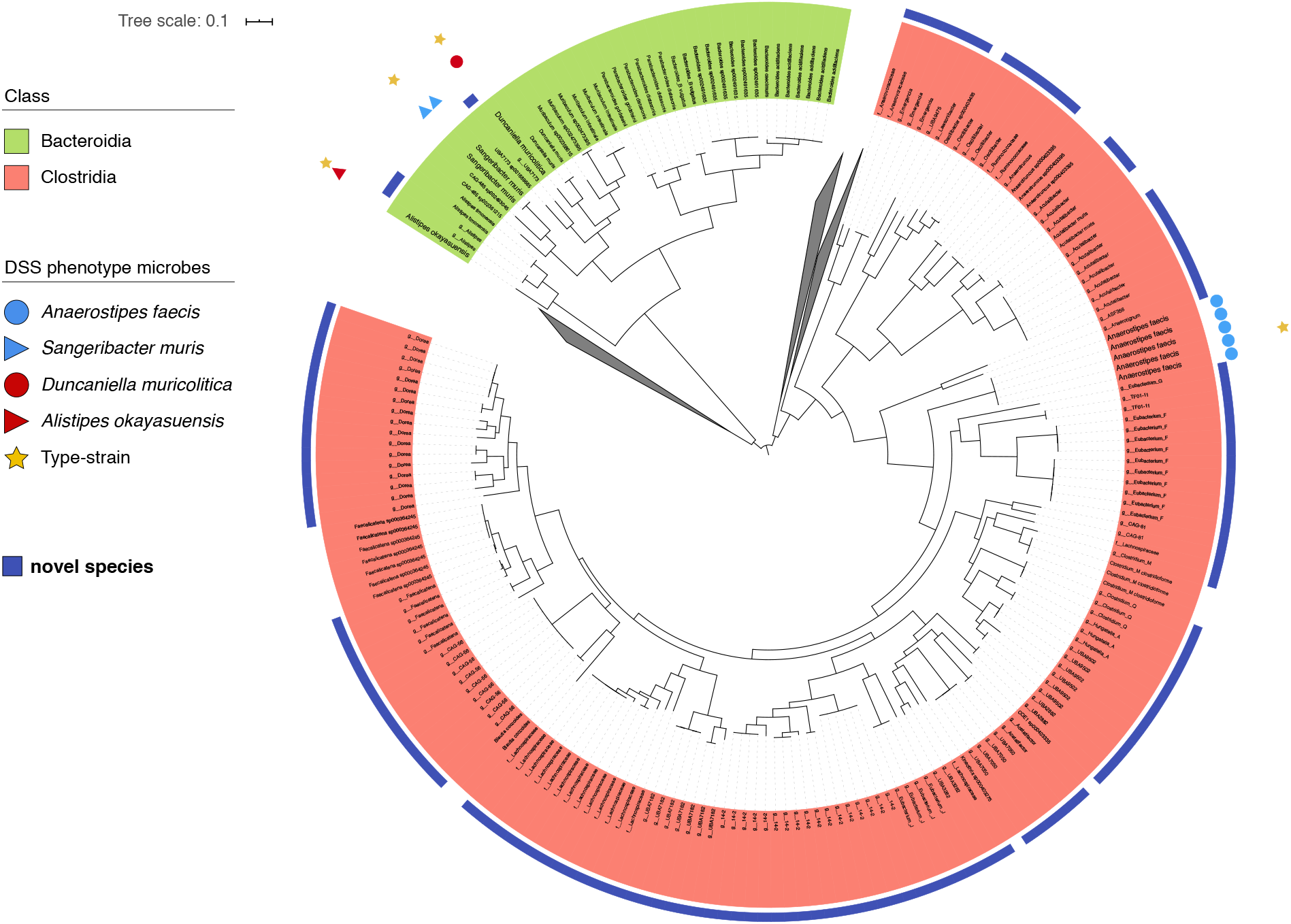
Genomic Characterisation of Bacterial Isolates. Phylogenetic tree showing relationship of *A. faecis* (blue circle), *S. muris* (blue triangle), *D. muricolitica* (red circle) and *A. okayasuensis* (red triangle) with known type strains (yellow star).

**Supplementary Figure 4:**
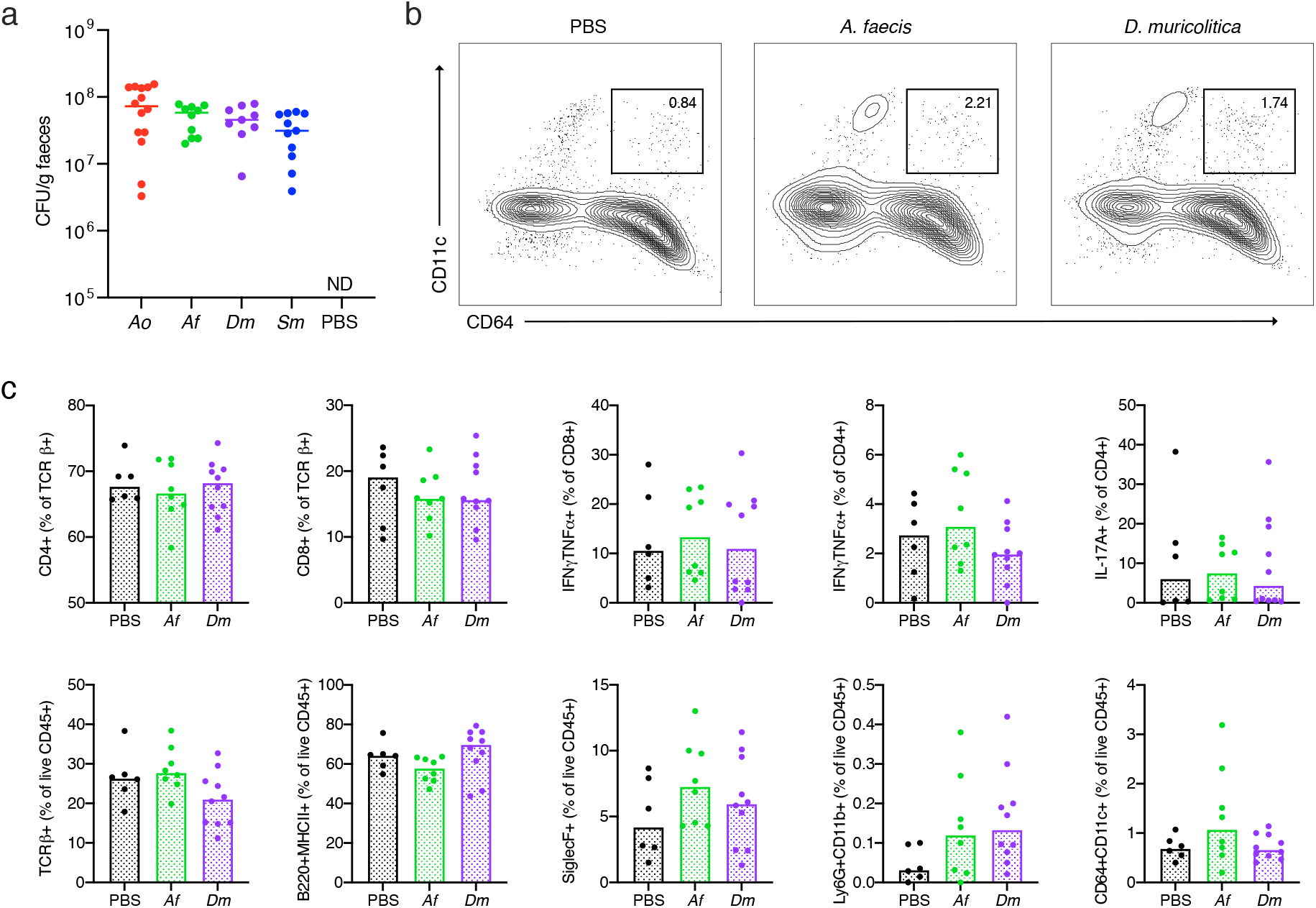
Mono-colonisation of germ-free mice. (a) Bacterial colonisation at day 0 prior to DSS treatment in mice pre-colonised with either *A. okayasuensis* (red), *A faecis* (green), *D. muricolitica* (purple) or *S. muris* (blue) and germ-free controls (PBS; black) presented as CFU/g of faecal material. No bacterial colonies were detected (ND) in the PBS control mice. Flow cytometric analysis of (b) monocytes/macrophages (B220/CD3e/TCRg-MHC2+CD64+CD11c+) and (c) other immune cell populations from the large intestine lamina propria of control germ-free mice (PBS) or mice mono-colonised with *A. faecis* or *D. muricolitica* without DSS challenge.

